# Raman spectral signatures reflect altered metabolic state of exhausted T cells

**DOI:** 10.64898/2026.06.16.732575

**Authors:** Ankita Arani Barai, Pooja C Asani, Partha Sarathi Mohanty, Arjit Tiwari, Sriya Bose, Soumen Das, Gayatri Mukherjee

## Abstract

T cell exhaustion within the tumor microenvironment drives CD8^+^ T cells into a dysfunctional state characterized by progressive loss of proliferative capacity and effector functions, thereby limiting anti-tumor immunity and therapeutic efficacy. To investigate the biochemical alterations associated with exhaustion, an in vitro model of CD8^+^ T cell exhaustion was established through chronic antigenic stimulation of murine OT-1 CD8^+^ T cells. Phenotypic, functional, metabolic, and transcriptional characterization confirmed the acquisition of an exhausted state. Single-cell Raman spectroscopy was subsequently employed to generate biochemical signatures of activated, and exhausted CD8^+^ T cells. Principal component analysis (PCA) of the Raman spectral data revealed distinct separation of these two cell subsets, reflecting underlying biochemical differences associated with their functional states. Differential Raman spectral features corresponding to nucleic acids, carbohydrates, proteins, and lipids contributed significantly to this segregation, reflecting altered metabolic and biosynthetic states during exhaustion progression. Classification of the spectral data using machine-learning algorithms enabled accurate segregation of activated and exhausted T cells. Collectively, this study demonstrates that single-cell Raman spectroscopy can distinguish exhausted CD8^+^ T cells in a label-free and non-destructive manner, highlighting its potential as a platform for immune profiling and monitoring T cell dysfunction.

## Introduction

Chronic antigenic stimulation within the tumor microenvironment (TME) drives effector CD8^+^ T cells into a state of exhaustion, characterized by progressive loss of proliferative capacity and cytotoxic function. This dysfunction compromises anti-tumor immunity, facilitating tumor progression and poor clinical outcomes. Under physiological conditions, naïve CD8^+^ T cells become activated upon recognition of tumor-associated or tumor-specific antigens (TAA/TSA) presented on major histocompatibility complex class I (MHC-I) molecules, followed by migration to the tumor site to eliminate malignant cells through cytotoxic granule release. However, persistent engagement of the T cell receptor (TCR) within the tumor milieu leads to sustained signaling, ultimately impairing effector function and rendering them exhausted.

Importantly, T cell exhaustion is not binary but exists along a continuum, with cells transitioning through distinct functional and phenotypic states before reaching terminal exhaustion marked by irreversible dysfunction. This continuum has been classified into stages based on the dynamics of CD69 and Ly108 expression, reflecting activation status and stemness^1^. Early exhausted populations include CD69^+^Ly108^+^ and CD69^−^Ly108^+^ subsets (Progenitor 1 and 2), which are more amenable to therapeutic reprogramming. As exhaustion progresses, Ly108 expression declines, leading to double-negative intermediate states and eventually terminally exhausted cells in which CD69 is re-expressed. These later stages show reduced responsiveness to immunotherapeutic interventions, with terminally exhausted cells being the most resistant.

Accurate identification of exhausted T cells within the TME is therefore critical for understanding disease progression and patient prognosis. The extent of exhausted CD8^+^ T cells can indicate tumor progression and help in selecting therapeutic strategies^2^. Consequently, there is a need for robust approaches to detect and characterize exhausted T cell populations in situ.

Exhausted T cells are typically identified through inhibitory receptor expression, transcriptional changes, and functional impairment^3^. These cells co-express PD-1, LAG-3, TIM-3, TIGIT, and CTLA-4^4^. Functionally, they exhibit reduced cytotoxicity due to decreased perforin and granzyme production^5^, along with diminished expression of the degranulation marker CD107a^6^. Cytokine production is also impaired, including IL-2^7^ and TNF-α^8^. Although IFN-γ is a proinflammatory cytokine that is often retained longer in exhausted cells, it can also exert regulatory effects through feedback mechanisms that limit anti-tumor immunity ^9^. Expression of the ectonucleotidase CD39 further marks terminally differentiated, irreversibly dysfunctional cells.

At the molecular level, exhaustion is driven by coordinated transcriptional and epigenetic reprogramming. Reduced TCF-1 and increased TOX expression are key features, with TOX acting as a central regulator that enforces the exhausted state^10^, while loss of TCF-1 promotes progression toward terminal exhaustion^11^. Additional transcription factors, including NFAT, BATF, and EOMES, further regulate exhaustion-associated gene programs^12^.

Current approaches for identifying exhausted T cells rely on surface marker profiling, functional assays, and transcriptional analyses. While widely used, these methods require extensive processing and may not fully capture the metabolic and biochemical complexity of exhaustion. Furthermore, overlap in inhibitory receptor expression between activated and transitional states limits their reliability in distinguishing functional phenotypes.

Given these limitations, there is a need for approaches that provide a direct, label-free readout of cellular biochemical states. In this context, Raman spectroscopy offers a useful alternative. Raman spectroscopy is a non-destructive, label-free technique that provides molecular-level information based on the inelastic scattering of light. When monochromatic laser light interacts with a sample, most photons undergo elastic (Rayleigh) scattering, while a small fraction interacts with molecular vibrations, resulting in energy shifts characteristic of Raman scattering. These shifts correspond to specific vibrational modes of chemical bonds and provide insight into molecular composition. By capturing vibrational signatures of lipids, proteins, and nucleic acids, Raman spectroscopy enables biochemical profiling at the single-cell level, generating intrinsic spectral fingerprints reflective of cellular states. Over the past decade, it has been applied in cancer diagnosis^13,14^ and in distinguishing cellular differentiation states^15,16,17^. More recently, it has been extended to immune cell characterization, including CD4^+^ T cells, B cells, and monocytes^18^, as well as differentiation between naïve and activated lymphocytes^19,20^. It has also been used to distinguish among T lymphocyte subsets, natural killer (NK) cells, and dendritic cells^21^, demonstrating its ability to resolve functional differences within immune populations. In particular, CD8^+^ T cells exhibit measurable spectral changes upon activation, supporting the reliability of this approach.

Beyond cellular characterization, Raman spectroscopy also holds diagnostic potential. For instance, metabolic profiling of NK cells has been used to distinguish cancer patients from healthy individuals and infer disease localization^22^. Given the central role of T cell exhaustion in tumor progression and its value as a prognostic biomarker^23^, identifying exhausted T cells based on intrinsic biochemical properties could provide significant clinical insight.

In the present study, we established an in vitro model of antigen-specific T cell exhaustion through chronic antigenic stimulation and performed phenotypic, functional, and transcriptional characterization. The system recapitulated a progressive exhaustion trajectory consistent with in vivo observations. Raman spectroscopy was used to acquire spectral profiles of exhausted cells and compared to those of activated CD8^+^ T cells. Principal Component Analysis of the Raman spectral data demonstrated segregation of exhausted and activated CD8^+^ T cells, indicating distinct biochemical states associated with exhaustion. Analysis of characteristic Raman bands revealed differences in peak intensities of wave-numbers corresponding to key biomolecular components, consistent with the metabolic and functional alterations observed in exhausted cells. Furthermore, classification of the spectral data using machine-learning algorithms enabled accurate discrimination of the different T-cell states. Collectively, these findings demonstrate that Raman spectroscopy can capture the biochemical landscape of CD8^+^ T cell exhaustion and represents a promising label-free approach for identifying exhausted T cells based on their intrinsic molecular signatures.

## Materials and Methods

### Mice and cell lines

OT-1 TCR Transgenic mice were obtained from The Jackson Laboratory, USA. Animals were housed in the Institutional Animal Facility under standard conditions with ad libitum access to food and water. All experiments were performed using male mice aged 10–12 weeks under the IAEC approval number IE-09/GM-SMST/1.22.

### Establishment of an in vitro reductionist model of T cell exhaustion

CD8+ T cells from the splenocytes of male OT-1 mice were isolated through negative selection using CD8^+^ T Cell Isolation Kit (BioLegend MojoSort™), according to the manufacturer’s protocol and cultured in complete RPMI 1640 medium supplemented with 10% FBS and 50 μM β-mercaptoethanol (R10). These cells were treated following the protocol mentioned below, to generate a reductionist in-vitro model of antigen-specific T cell exhaustion.

For generating acutely stimulated (AS) as well as chronically stimulated (CS) cells, initially the CD8+ T cells were seeded at a density of 0.2 x 10^6 cells/well on 96 well plates coated with 1ug/ml of SIINFEKL-□2m-H2Kb pMHC complex, produced in-house according to previously published protocol (Mahata et al., 2023). After 48 hours, cells for the AS set were harvested, washed and reseeded at the same density in fresh media without any pMHC stimulation. For the CS sets, at every 48 hour interval, the cells were harvested, counted and reseeded with pMHC stimulation. The first restimulation was given on day 2 with plate-bound pMHC, and the cells were kept in conditioned media (CM)., The subsequent restimulations on day 4 and 6 were given with pMHC in a soluble form and the cells were cultured in 1: 1 CM: fresh R10 at every time point. This sequence of stimulation generated a continuum of exhaustion stages in these cells at day 4, 6 & 8. Untreated cells on day0 were seeded in the same complete RPMI 1640 medium without stimulation and kept as negative control.

The cells were harvested at specific time points for flow cytometry and Raman spectroscopy and cell supernatant was collected for performing ELISA for different cytokines. For all the experiments splenocytes from more than two mice were pooled together and multiple biological replicates (3-5) were used.

### Assessment of T Cell Proliferation

The proliferative capacity of CD8^+^ T cells during the course of exhaustion was assessed by calculating population doublings at each stimulation time point. Cells were harvested and counted every 48 h, corresponding to each round of stimulation. Population doublings (PD) were calculated by comparing the number of recovered cells (N_f_) at a given time point with the initial number of cells seeded (N_i_) according to the following equation:

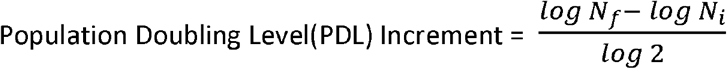

### Flow Cytometry

The phenotypic and functional conditions of the exhausted cells were investigated by multi-colour Flow cytometry. Briefly, at each time point, the cells were harvested, counted and stained with the following flurochrome-conjugated monoclonal antibodies (mAbs): anti-CD8 FITC (53-6.7) (BD Biosciences), anti-CD8 PercPCy5.5 (53-6.7) (BD Biosciences), anti-CD69 PECF594 (H1.2F3) (BD Biosciences), anti-Ly-108 APC (330-AJ) (BioLegend), anti-PD-1 APC (J43) (BD Bioscience), anti-PD-1 PE (J43) (BD Bioscience), anti-TIM-3 BV421 (RMT3-23) (BD Bioscience), anti-TIGIT PE (1G9) (BD Bioscience), anti-LAG-3 PE-Cy7 (C9B7W) (Invitrogen), and anti-CTLA-4 PE (UC10-4F10-11) (BD Bioscience), anti-granzyme-B PE-Cy7 (QA18A28) (BioLegend), anti-perforin Pacific Blue (S16009A) (BioLegend), anti-CD39 BV421 (Y23-1185) (BD Bioscience), anti-CD38 BV650 (90/CD38) (BD Bioscience), anti-CD36 PECy7 (HM36) (BioLegend).

Before staining the cells for surface or intra-cellular markers, all cells were stained with BD Horizon™ Fixable Viability Stain 510, to enumerate cell viability and also to ensure the selection of live cells only for flow cytometric analysis. Then the cells were washed with FACS buffer (1% FBS in 1X PBS pH 7.2-7.4) and incubated with Fc block™ (clone: 2.4G2) (BD Biosciences) for 5-10 minutes in order to avoid non-specific binding of antibodies through their Fc regions. Cells were washed and subsequently stained with the flurochrome-conjugated mAbs specific for the different surface markers. Cells were washed with FACS buffer and collected in FACS buffer for data acquisition.

For intracellular staining of Granzyme, Perforin, and cytokines, the AS & CS cells were given an extra stimulation of pMHC (1ug/ml) along with BD GolgiPlug™ Protein Transport Inhibitor (Containing Brefeldin A) and incubated for 6 hours, at 37dC, 5% CO2, before the cells were harvested for staining. For intracellular staining of transcription factors (TOX & TCF), no such stimulation or golgi plug was required. After the surface staining, the cells were fixed for 20 mins on ice and washed with 1X Permwash using the BD Cytofix/Cytoperm™ Fixation/Permeabilization Kit. The cells were then stained with the intracellular antibodies diluted in the same buffer by incubating them for 30 minutes. The cells were washed with FACS buffer and collected as mentioned earlier for data acquisition.

For the degranulation marker, CD107a, cells were similarly re-stimulated with pMHC complex, in presence of anti-CD107a antibody (ID4B) (BD Biosciences), followed by BD GolgiStop™ after 1 hour and incubated for a total of 6 hours at 37°C, 5% CO2., according to manufacturer’s instructions. Cells were then stained with surface markers as required and collected for acquisition.

Data from these samples were acquired on a LSRFortessa (BD Biosciences) and analysed using FlowJo™ software.

### ELISA

Cell-free supernatants collected from chronically stimulated (CS) and acutely stimulated (AS) CD8^+^ T cells were analyzed for cytokine secretion. Levels of IL-2 were quantified using mouse DuoSet ELISA kit (R&D Systems) according to the manufacturer’s instructions. Briefly, 96-well plates were coated overnight with capture antibodies, followed by blocking with 1% BSA in 1X PBS. Samples and standards were then added and incubated, followed by detection antibodies and streptavidin–HRP. Signal was developed using substrate solution, and the reaction was stopped with stop solution. Absorbance was measured at 450 nm & 540nm (for background subtraction) using a microplate reader, and cytokine concentrations were determined from standard curves.

### Confocal Raman Spectroscopy of CD8^+^ T cells subsets

48 h-activated T cells, and chronically stimulated T cells at day 6 (CS6) were harvested in each experimental round and fixed with 4% paraformaldehyde for 30 minutes. Following fixation, cells were washed twice with 1× PBS to remove residual fixative, resuspended in PBS, and drop-cast onto sterile, negatively charged glass slides. The slides were air-dried at room temperature prior to Raman measurements.

Single-cell Raman spectra were acquired using an Oxford WiTec UHTS 300 system equipped with an alpha300 RA microscope. Cells were visualized under a 50× air objective, and individual cells were probed using a 532 nm excitation laser. Spectral acquisition was performed using WiTec Control 7.0 software, and baseline correction was carried out in WiTec Project 7.0.

All spectra were recorded with a center wavelength of 602.593 nm and a 600 g/mm diffraction grating. Each spectrum was obtained as an average of 10 accumulations with an integration time of 5 s per accumulation, optimized to ensure spectral consistency. Single-cell spectra were collected from randomly selected fields for each T cell subset across three independent biological replicates, with a total of 120 spectra acquired per subset.

### Data analysis of the Raman spectral signatures

The raw Raman spectra were pre-processed in order to improve the signal quality before proceeding for analysis. High-intensity cosmic ray artifacts were removed using the Cosmic Ray Removal (CRR) function implemented in the acquisition software. Baseline correction was performed using WiTec Project 7.0, and the corrected spectra were exported for further processing.

Following pre-processing, the data was plotted as intensity versus wavenumber in order to generate representative spectral profiles for each cell subset. Data processing was performed using Origin 2024.

The pre-processed spectral data from individual cells were imported into RStudio for performing multivariate analysis. The data matrix was organized with wavenumbers as variables and individual spectra as observations. Multivariate analysis was carried out using Principal Component Analysis (PCA) in RStudio. PCA was done using the prcomp function from the stats package, which applies singular value decomposition (SVD) to a mean-centered dataset. Unit variance scaling was done to standardise the contribution from each wavenumber, which ultimately helps effectively perform the PCA on correlation matrix. PCA scores and loadings were extracted and used for visualization and interpretation of spectral patterns. Significant peaks were selected for different biomolecules and analysed for area under the curve using Origin 2024 & GraphPad Prism.

### Raman Spectral Data Processing and Machine Learning Classification

The pre-processed Raman spectra were organized into a feature matrix in which each row represented an individual cell spectrum and each column corresponded to the Raman intensity at a specific wavenumber. The final dataset consisted of 240 spectra and 1650 spectral features. Activated and exhausted T-cell spectra were assigned binary class labels for supervised classification.

Prior to analysis, spectral features were standardized using z-score normalization according to the equation:

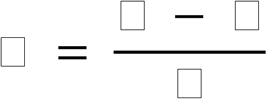

where (x) is the original feature value, (µ) is the mean feature value, and (σ) is the corresponding standard deviation. Standardization was performed to minimize scale-dependent variation among spectral variables and improve model performance.

To evaluate the ability of Raman spectral signatures to discriminate T-cell states, three supervised machine learning algorithms were implemented: Random Forest (RF), Support Vector Machine (SVM), and Extreme Gradient Boosting (XGBoost). RF was constructed using 500 decision trees, SVM was implemented with a radial basis function (RBF) kernel, and XGBoost was trained using 300 estimators with a maximum tree depth of 4 and a learning rate of 0.05. In addition, Principal Component Analysis (PCA) was applied as a dimensionality reduction strategy, and classifier performance was assessed using both the original spectral features and the PCA-transformed feature space.

### Model Evaluation and External Validation

Classifier performance was evaluated using stratified five-fold cross-validation to ensure proportional representation of activated and exhausted T-cell spectra within each fold. During each iteration, four folds were used for training and one-fold for validation, such that every sample was evaluated once. Performance was assessed using classification accuracy, receiver operating characteristic (ROC) analysis, area under the ROC curve (AUC), confusion matrix, precision, recall, and F1-score. Mean accuracy and standard deviation across the five folds were calculated to assess model robustness and generalizability.

The performances of RF, SVM, and XGBoost were compared under identical evaluation conditions to identify the most suitable classifier for Raman-based discrimination of T-cell states. The best-performing model was subsequently selected for detailed ROC and confusion matrix analyses.

To assess model generalizability, external validation was performed using an independent Raman spectral dataset acquired from biologically distinct samples that were not included in model training or cross-validation. The validation dataset consisted of 40 activated-cell spectra and 40 exhausted-cell spectra. The trained XGBoost classifier was applied directly to the external dataset without retraining. Classification performance was evaluated using accuracy, confusion matrix analysis, precision, recall, F1-score, and AUC, enabling assessment of model robustness across independent exhausted T-cell populations.

### Statistical analysis

All experimental data were verified by multiple biological replicates. GraphPad Prism (v9) was used to perform statistical analysis. p-value was calculated using one-way ANOVA and t Test wherever applicable and statistical significance was indicated as *p < 0.05, **p < 0.01, ***p < 0.001, and ****p < 0.0001.

## Results

### Generation of an in-vitro exhaustion model

An in vitro model of T cell exhaustion was established by chronically stimulating OT-1 CD8^+^ T cells with SIINFEKL-H2K⍰ complexes. As illustrated schematically in Fig. 1A, cells were stimulated every 48 hours until day 8. The initial two stimulations were performed using plate-bound pMHC complexes, following which the cells were maintained in conditioned media until 96 hours. Subsequent stimulations were carried out in soluble form with half-media replacement using fresh media. As controls, cells were either maintained without stimulation (UT; untreated) or rested following the initial stimulation to generate acutely stimulated (AS) cells. Under these conditions, chronically stimulated (CS) cells progressively acquired phenotypic, transcriptional, metabolic and functional features associated with T cell exhaustion.

**Figure 1.**
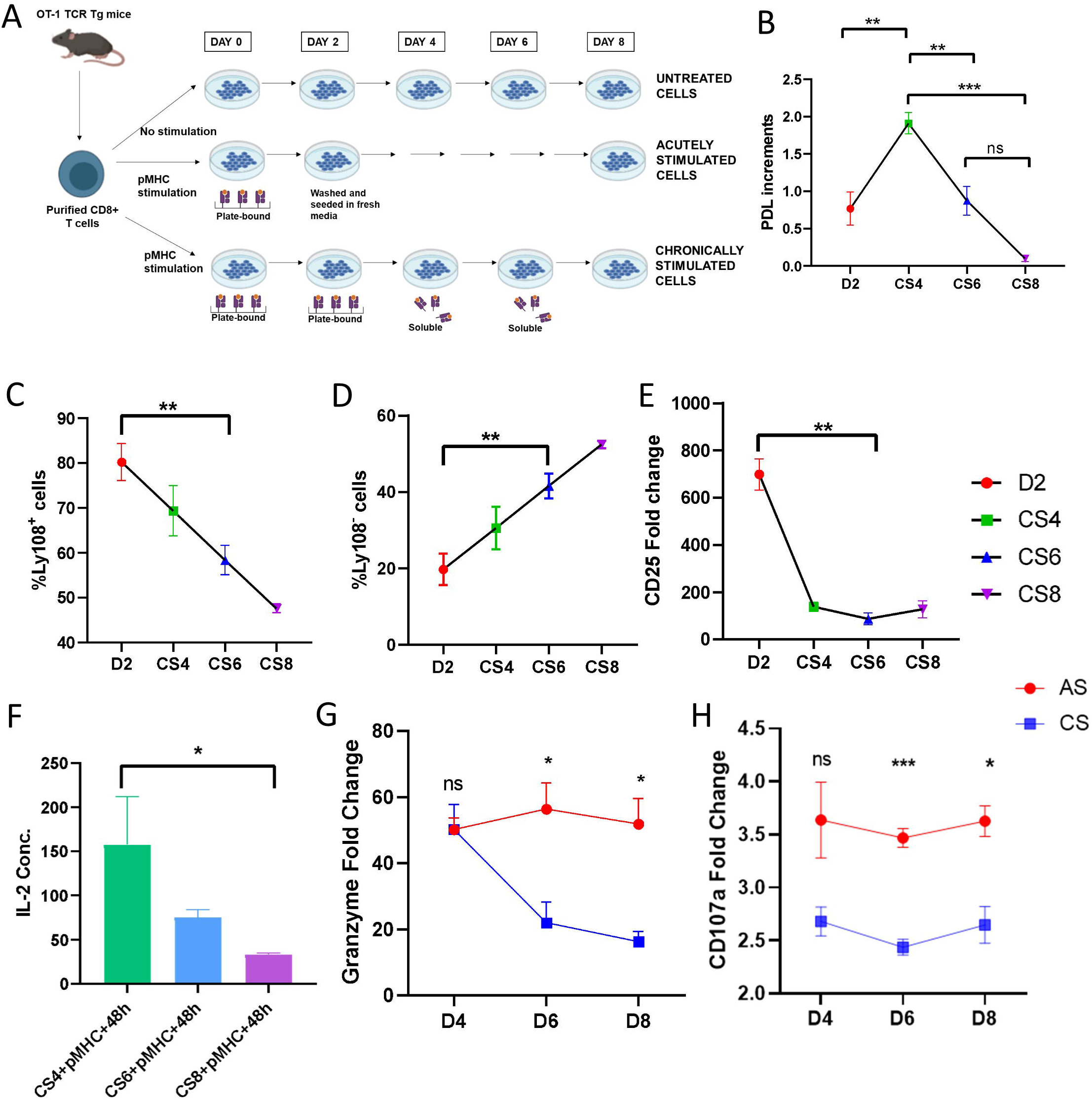
Establishment of an in vitro CD8^+^ T cell exhaustion model associated with progressive loss of effector function. (A) Schematic representation of the experimental workflow used for the generation of the in vitro exhaustion model through repeated antigenic stimulation. (B) Population doubling levels of CS cells across different stages of exhaustion. (C) Percentage of Ly108^+^ progenitor-like exhausted cells and (D) Ly108^−^ intermediate/terminally exhausted cells at the indicated timepoints. (E) Quantification of CD25 expression during exhaustion progression. (F) IL-2 concentration in culture supernatants collected from cells at 48 hours post repeat stimulation with pMHC on days 4, 6, and 8. (G) Granzyme expression and (H) CD107a degranulation levels in CS cells compared to AS controls at the indicated timepoints. Data are represented as mean ± SEM from independent biological replicates (n=3-6). Statistical significance was determined using unpaired Student’s t-test.

CS cells exhibited a progressive decline in proliferative capacity with repeated stimulation (Fig. 1B). A proliferative burst was observed around day 4, suggestive of a progenitor-like exhausted state, following which proliferation gradually declined.

To further assess exhaustion progression, the distribution of progenitor exhausted (Ly108^+^) and intermediate/terminally exhausted (Ly108^−^) populations was examined across different timepoints. CS cells displayed a gradual reduction in the Ly108^+^ population (Fig. 1C) accompanied by a steady increase in Ly108^−^ cells (Fig. 1D), indicating progressive differentiation toward more exhausted states. Consistent with this transition, CD25 expression was significantly reduced (Fig. 1E), along with a corresponding decrease in IL-2 production (Fig. 1F). In agreement with these observations, chronically stimulated cells also exhibited marked functional impairment compared to acutely stimulated cells, as evidenced by reduced Granzyme (Fig. 1G) and CD107a expression (Fig. 1H). Collectively, these findings confirmed the successful establishment of a progressive in vitro T cell exhaustion model.

### Phenotypic characterization of CS6 cells

Chronically stimulated day 6 (CS6) cells exhibited a high proportion of intermediately exhausted cells, accompanied by significant loss of proliferative capacity and effector function. Therefore, these cells were further characterized for the expression of exhaustion-associated inhibitory receptors.

Within the tumor microenvironment, exhausted T cells undergo functional suppression through sustained engagement of inhibitory receptors. Consequently, co-expression of multiple inhibitory receptors serves as an important hallmark of T cell exhaustion. Both AS and CS cells displayed elevated PD-1 expression (Fig. 2A), although more pronounced differences were observed for LAG-3 expression, where a substantial population of CS cells co-expressed PD-1 and LAG-3, while such populations were nearly absent in AS cells (Fig. 2B).

**Figure 2.**
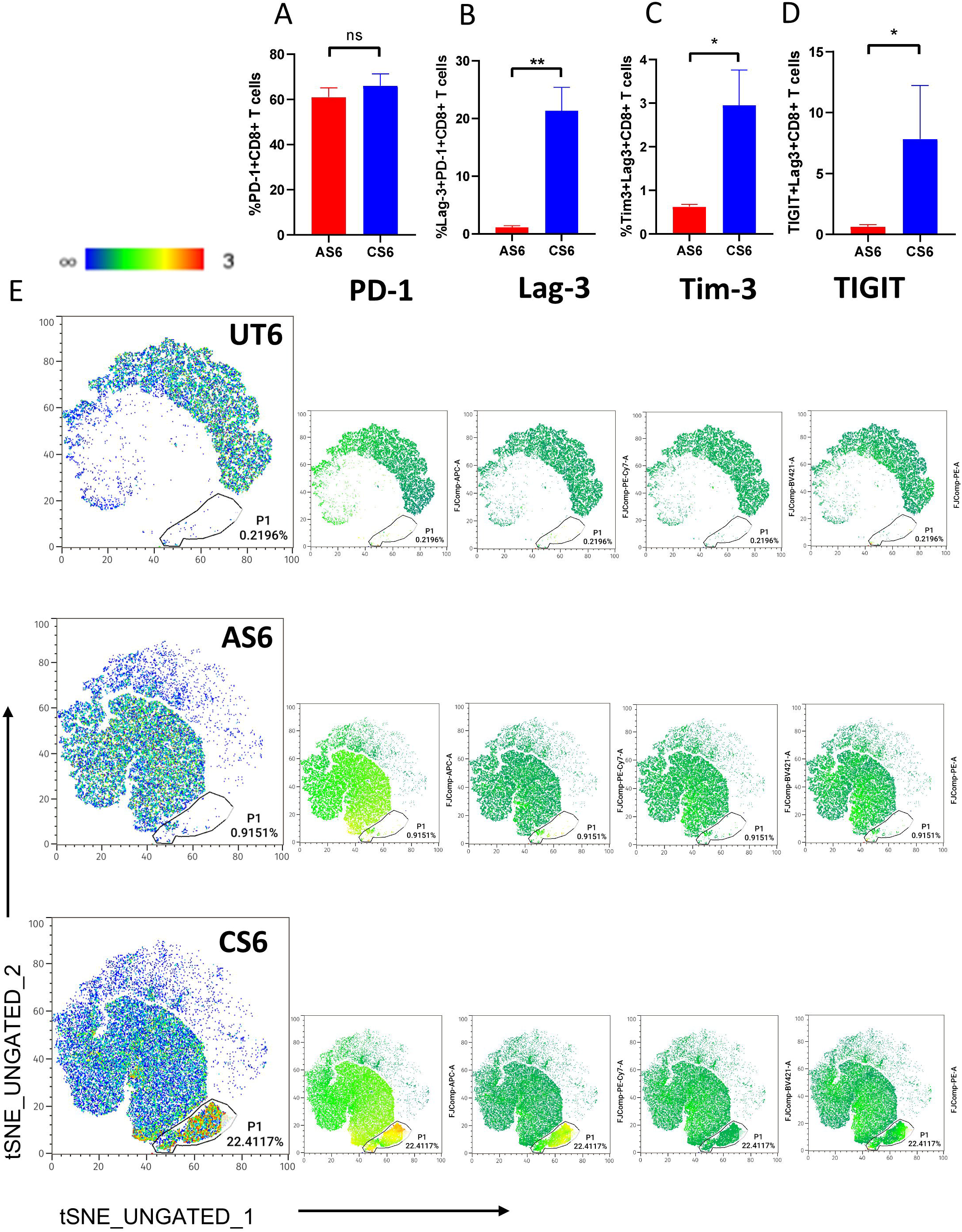
Phenotypic characterization of chronically stimulated CD8^+^ T cells. (A) Percentage of PD-1^+^ cells, (B) Percentage of LAG-3^+^PD-1^+^ cells, (C) Percentage of Tim3^+^LAG-3^+^PD-1^+^ cells, (D) Percentage of TIGIT^+^LAG-3^+^PD-1^+^ cells in AS & CS sets. (E) tSNE clustering demonstrating co-expression patterns of PD-1, LAG-3, TIM-3 & TIGIT within AS & CS cells. Data are represented as mean ± SEM from independent biological replicates (n=3). Statistical significance was determined using unpaired Student’s t-test.

Along with Lag-3, CS6 cells also showed higher expression of other inhibitory receptors like TIGIT & Tim-3 in comparison to AS cells (Fig. 2C & 2D). Furthermore, chronically stimulated cells demonstrated increased co-expression of exhaustion-associated inhibitory receptors, as visualized through t-SNE analysis (Fig. 2E). The distinct clustered population observed exclusively within the CS cells, but absent in AS cells, was characterized by high PD-1 and LAG-3 expression with moderate TIGIT and TIM-3 expression, further supporting the acquisition of an exhaustion-associated phenotype.

### Metabolic and transcriptional characterisation of CS6 exhausted cells

T cell exhaustion is associated not only with inhibitory receptor expression and functional decline, but also with extensive metabolic and transcriptional reprogramming. In the tumor microenvironment, exhausted T cells are often characterized by altered lipid metabolism and increased accumulation of fatty acids, which contribute to cellular dysfunction. Consistent with this phenotype, chronically stimulated day 6 (CS6) cells exhibited elevated expression of the lipid scavenger receptor CD36 compared to acutely stimulated (AS6) cells, suggesting enhanced lipid uptake and metabolic dysregulation (Fig. 3A). In addition, CS6 cells showed increased expression of CD39, an ectonucleotidase associated with terminal exhaustion and immunosuppressive T cell states (Fig. 3B).

**Figure 3.**
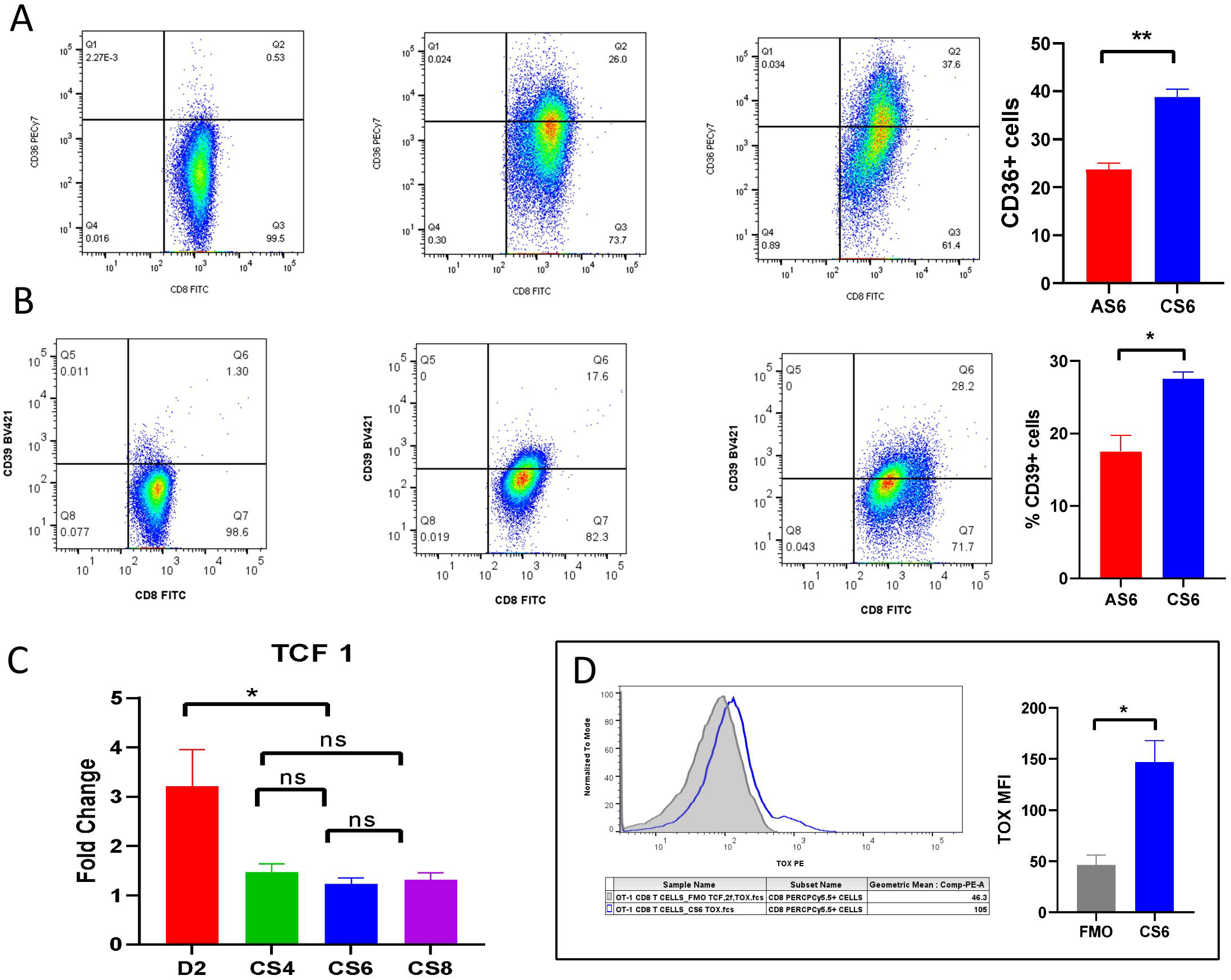
Metabolic and transcriptional characterization of chronically stimulated CD8^+^ T cells on day 6. (A) Representative flow cytometry dot plots showing CD36 expression in UT, AS6, and CS6 cells, along with pooled quantification of the percentage of CD36^+^ cells in AS6 and CS6 populations. (B) Representative flow cytometry dot plots showing CD39 expression in UT6, AS6, and CS6 cells, with pooled quantification of the percentage of CD39^+^ cells. (C) Progressive loss of TCF-1 transcription factor from day 2 to day 8 (D) Representative histogram plots showing TOX expression relative to FMO controls in CS6 cells, with corresponding pooled MFI quantification. Data are represented as mean ± SEM from independent biological replicates (n=3). Statistical significance was determined using an unpaired Student’s t-test.

Exhausted T cells also undergo characteristic transcriptional changes that regulate and sustain the dysfunctional phenotype. Among these, TOX and TCF-1 are key transcriptional regulators associated with exhaustion progression. The exhausted cells displayed markedly reduced TCF-1 expression, as they progress towards more terminal stages, indicating loss of stem-like and progenitor characteristics (Fig. 3C). The CS6 exhausted cells which were nearly negative for TCF-1, were found to have sustained expression of TOX (Fig. 3D), supporting the acquisition and maintenance of an exhaustion-associated transcriptional program.

### Raman spectral signatures produce distinct clustering of activated & exhausted cells

To investigate whether T cell exhaustion can be discriminated from activated CD8+ T cells through non-invasive ways, single-cell Raman spectra were acquired from activated, and chronically stimulated day 6 (CS6) CD8^+^ T cells. Comparative analysis of the averaged spectra revealed differences in many peak intensities across the three cellular states, indicating underlying biochemical variations associated with activation and exhaustion. Raman spectra acquired from the cells were initially visualized across the full spectral range (Fig. 4A). For detailed biochemical interpretation, subsequent analysis was focused on the fingerprint region between 250–1800 cm^−1^, which encompasses the major biologically relevant Raman bands while excluding the Raman silent region (Fig. 4C).

**Figure 4.**
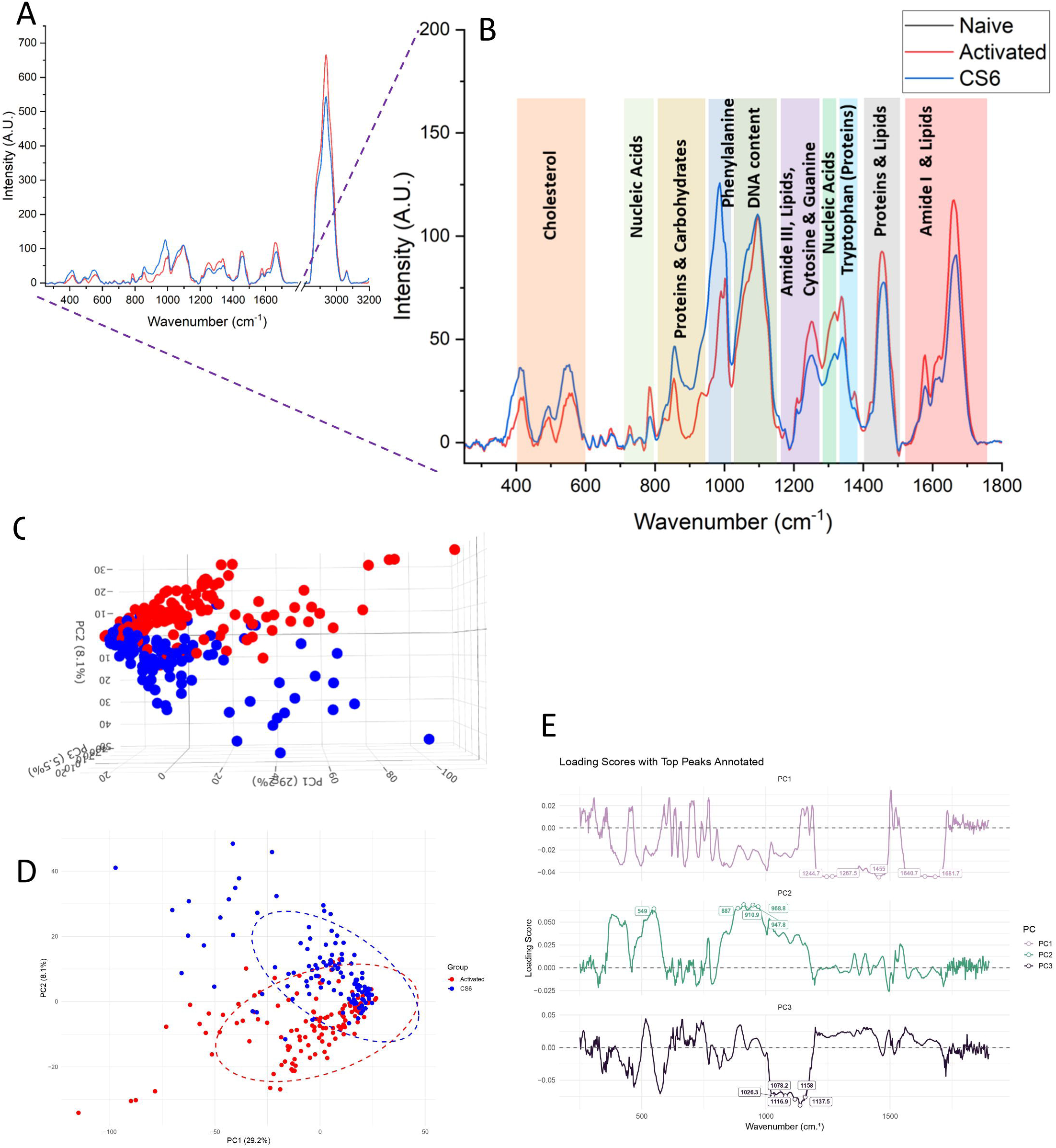
Raman spectroscopic profiling of activated, and exhausted CD8^+^ T cells. (A) Averaged Raman spectra of activated, and CS6 CD8^+^ T cells across the full spectral range (120 cells for each set from 3 independent biological replicates). (B) Expanded view of the biologically relevant fingerprint region (250–1800 cm^−1^) highlighting Raman peaks corresponding to major biomolecular vibrations. (C) Principal Component Analysis (PCA) 3D plot & (D) 2D convex hull demonstrating clustering and spectral segregation of activated, and CS6 CD8^+^ T cell populations.(E) Loading Scores with Top Peak Annotations in PC1, PC2 & PC3

The analysis revealed distinct biochemical alterations across activated and exhausted CD8^+^ T cell states. To further evaluate whether these spectral differences could distinguish the cellular states in an unbiased manner, Principal Component Analysis (PCA) was performed using single-cell Raman spectra of activated and exhausted (CS6) cells. PCA demonstrated distinct clustering of activated and exhausted T cell populations in both two-dimensional and three-dimensional score plots (Fig. 4D). Activated cells and exhausted cells localized in separate clusters, indicating partial loss of the biochemical characteristics associated with T cell activation. This clustering pattern supports the progressive functional and metabolic decline observed during exhaustion. To identify the spectral regions contributing to the separation of the cell populations, loading scores for principal components 1, 2, and 3 were examined. The major discriminating wavenumbers identified from the loading plots are summarized in Table 1.

**Table 1:** Loading scores for PC1, PC2 & PC3.

| PC | Rank | Wavenumber | Loading | AbsLoading | Direction |
| --- | --- | --- | --- | --- | --- |
| PC1 | 1 | 1455.034 | -0.04445 | 0.044453 | Negative |
| PC1 | 2 | 1452.562 | -0.04445 | 0.044446 | Negative |
| PC1 | 3 | 1457.505 | -0.0443 | 0.044304 | Negative |
| PC1 | 4 | 1450.09 | -0.0442 | 0.044202 | Negative |
| PC1 | 5 | 1681.658 | -0.04411 | 0.044106 | Negative |
| PC1 | 6 | 1679.254 | -0.0441 | 0.044095 | Negative |
| PC1 | 7 | 1244.737 | -0.04408 | 0.044079 | Negative |
| PC1 | 8 | 1249.808 | -0.04405 | 0.044052 | Negative |
| PC1 | 9 | 1684.06 | -0.04405 | 0.044049 | Negative |
| PC1 | 10 | 1676.851 | -0.04402 | 0.044023 | Negative |
| PC2 | 1 | 910.8561 | 0.069127 | 0.069127 | Positive |
| PC2 | 2 | 947.7559 | 0.068668 | 0.068668 | Positive |
| PC2 | 3 | 908.2141 | 0.068245 | 0.068245 | Positive |
| PC2 | 4 | 945.1257 | 0.06813 | 0.06813 | Positive |
| PC2 | 5 | 950.3852 | 0.068119 | 0.068119 | Positive |
| PC2 | 6 | 953.0137 | 0.068035 | 0.068035 | Positive |
| PC2 | 7 | 913.4974 | 0.067821 | 0.067821 | Positive |
| PC2 | 8 | 955.6414 | 0.067608 | 0.067608 | Positive |
| PC2 | 9 | 960.8942 | 0.067529 | 0.067529 | Positive |
| PC2 | 10 | 905.5712 | 0.067436 | 0.067436 | Positive |
| PC3 | 1 | 1137.518 | -0.08848 | 0.088485 | Negative |
| PC3 | 2 | 1134.948 | -0.08656 | 0.086563 | Negative |
| PC3 | 3 | 1140.087 | -0.08625 | 0.086255 | Negative |
| PC3 | 4 | 1142.656 | -0.08514 | 0.085138 | Negative |
| PC3 | 5 | 1145.223 | -0.08238 | 0.082381 | Negative |
| PC3 | 6 | 1147.79 | -0.08188 | 0.081882 | Negative |
| PC3 | 7 | 1132.377 | -0.08143 | 0.081426 | Negative |
| PC3 | 8 | 1152.921 | -0.08083 | 0.08083 | Negative |
| PC3 | 9 | 1155.486 | -0.08039 | 0.080395 | Negative |
| PC3 | 10 | 1150.356 | -0.07998 | 0.07998 | Negative |

### Quantitative analysis of Raman bands reveals biomolecular features distinguishing exhausted and activated CD8^+^ T cells

To determine whether Raman spectroscopy could capture biomolecular differences associated with T cell functional states, selected Raman bands identified from the PCA loading scores, along with additional bands corresponding to biologically relevant molecules, were quantitatively analyzed using area under the curve (AUC) measurements. The Raman bands at ~426 cm^−1^ and ~546 cm^−1^, corresponding to the skeletal vibration and ring-breathing modes of cholesterol, respectively^24^, showed significantly higher AUC values in exhausted cells compared with activated cells (Fig. 5A–B). This increase suggests an accumulation of cholesterol in exhausted cells, potentially resulting from enhanced lipid uptake together with impaired cholesterol efflux. Such observations are consistent with reports demonstrating dysregulation of cholesterol transporters such as ABCA1 and ABCG1, which mediate cholesterol export from T cells^25^, as well as increased expression of the lipid scavenger receptor CD36, which facilitates uptake of oxidized LDL and cholesterol-containing lipids^26^.

**Figure 5.**
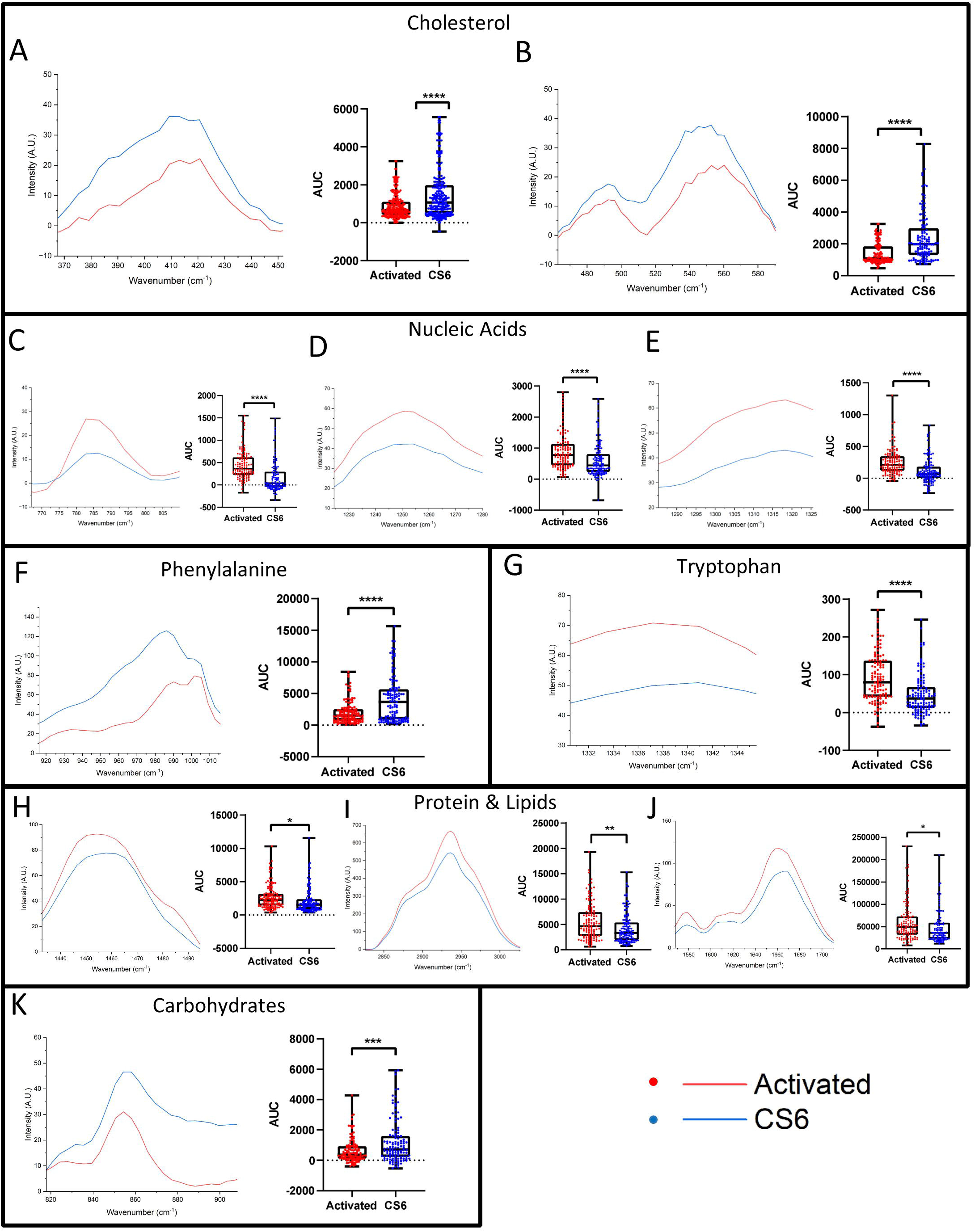
Quantitative analysis of biomolecule-associated Raman spectral features in activated and exhausted CD8^+^ T cells. Averaged Raman spectra and corresponding area under the curve (AUC) quantification of individual-cell spectra for cholesterol-associated bands (A-B), Nucleic Acid-associated bands (C-E), Phenylalanine-associated band (F), Tryptophan-associated band (G), Protein & Lipid-associated bands (H-J) & Carbohydrate-associated band (K) for activated and exhausted cells

Analysis of nucleic acid-associated Raman bands revealed distinct differences between activated and exhausted cells. Bands around ~780 cm^−1^, attributed to cytosine ring-breathing vibrations, ~1250 cm^−1^, associated with uracil and cytosine ring stretching, and ~1320 cm^−1^, corresponding to adenine and guanine ring stretching vibrations^27^, exhibited higher intensities in activated cells. These findings are consistent with the increased proliferative activity and elevated rates of transcription and nucleic acid synthesis that accompany T cell activation.

A prominent difference was also observed in the phenylalanine-associated band at 1003–1005 cm^−1^, which corresponds to the symmetric ring-breathing vibration of the phenyl group^28^. Exhausted cells displayed significantly higher intensity in this region compared with activated cells. This enhanced signal may reflect altered amino acid metabolism and increased intracellular phenylalanine levels. Such metabolic alterations are compatible with the bioenergetic dysfunction characteristic of exhausted T cells, where impaired mitochondrial fitness and oxidative phosphorylation can disrupt nutrient utilization. Elevated phenylalanine has been reported to suppress proliferation of CD4^+^ T cells through modulation of STAT6 and mTOR signaling pathways^29^. Given the importance of these pathways in CD8+ T cell activation and function, the increased phenylalanine-associated signature observed here may represent a metabolic feature associated with the diminished proliferative and effector capacity of exhausted cells.

In contrast, the tryptophan-associated band at ~1340 cm^−1 28^ showed significantly lower intensity in exhausted compared with activated cells. This reduction suggests alterations in tryptophan metabolism during exhaustion and may be indicative of enhanced catabolism through pathways such as the kynurenine pathway, which has been implicated in the regulation of immunosuppressive gene programs^30^.

Several protein- and lipid-associated Raman bands also differed between activated and exhausted cells. The band around ~1440 cm^−1^, arising primarily from CH_2_ deformations in lipid acyl chains and CH_3_ bending vibrations of aliphatic amino acid side chains^31^, showed reduced intensity in exhausted cells relative to activated cells. Similarly, the broad spectral region between 2800–3000 cm^−1^, dominated by CH_2_ and CH_3_ stretching vibrations from proteins and lipids^31^, exhibited higher intensity in activated cells. Additional differences were observed in the amide-associated regions, including the band around ~1580 cm^−1^, which encompasses contributions from Amide II vibrations involving N–H bending and C–N stretching^31^, and the region near ~1650 cm^−1^ corresponding to the Amide I band, dominated by C=O stretching of peptide backbones and C=C stretching of unsaturated fatty acids^31^. Collectively, these observations indicate increased protein synthesis and lipid-associated biochemical activity in activated cells, consistent with their heightened metabolic and effector functions.

Finally, exhausted cells exhibited relatively higher intensities within the carbohydrate-associated spectral region (~850–950 cm^−1^) compared with activated cells. This region has previously been associated with glycogen and carbohydrate-related vibrational signatures^32^. Several studies have demonstrated that exhausted T cells exhibit aberrant glycosylation, especially in the form of β1,6-GlcNAc-branched N-glycans and manipulating the glycosylation pathway offers a therapeutic strategy of reprogramming exhausted T cells^33,34^. The observed increase in carbohydrate-related peak intensities therefore could be reflective of the changes in glycosylation patterns in the exhausted T cells and warrants further in-depth studies. Further, while most systems of T cell exhaustion report a decrease in downregulation of GLUT1 receptor and glucose metabolism in T cell exhaustion^35^, one study with exhausted T cell in chronic viral infection showed upregulation of glucose uptake pathways accompanied by overexpression of the Glut1 glucose transporter^36^. TTherefore, the elevated carbohydrate-associated Raman signal may also indicate increased glucose uptake, albeit resulting in altered carbohydrate utilization and metabolic reprogramming rather than efficient glucose catabolism. Together, these findings demonstrate that Raman spectroscopy captures distinct biomolecular changes associated with T cell activation and exhaustion, providing a label-free readout of their underlying metabolic and functional states.

### Machine Learning Classification of T Cell States Based on Raman Spectra

To evaluate the discriminatory efficiency of Raman spectral signatures, Random Forest (RF), Support Vector Machine (SVM), and Extreme Gradient Boosting (XGBoost) classifiers were trained using stratified five-fold cross-validation, on the training dataset.. ROC analysis demonstrated strong classification performance, with all models achieving AUC values greater than 0.9, and RF showing the highest AUC (0.941) (Fig. 6B). But, when looking at model accuracy in Fig. 6A and their respective standard deviations, XGBoost came out to be the best classifier with accuracy (85.8 ± 3.58%), followed by RF (85.4 ± 4.16%) and SVM (83.8 ± 4.82%). Dimensionality reduction was performed using PCA along with these classifiers, but it did not improve the model performances, thus indicating that the Raman spectra has sufficient discriminatory information for classification. Therefore, based on its overall performance, XGBoost was selected for further evaluation. When trained with 80% of the data, it achieved an accuracy of 83.3% with only eight misclassified spectra in the 20% test set (Fig. 6C). For an external validation, independent samples were also run for testing, where it demonstrated robust classification performance, achieving 100% accuracy for all the activated and exhausted cells with an AUC of 1.0 (Fig. 6D). Collectively, these findings demonstrate that Raman spectral fingerprints contain sufficient biochemical information to enable accurate machine learning-based discrimination of activated and exhausted CD8□ T cells, supporting the potential of Raman spectroscopy as a label-free platform for immune cell state classification.

**Figure 6.**
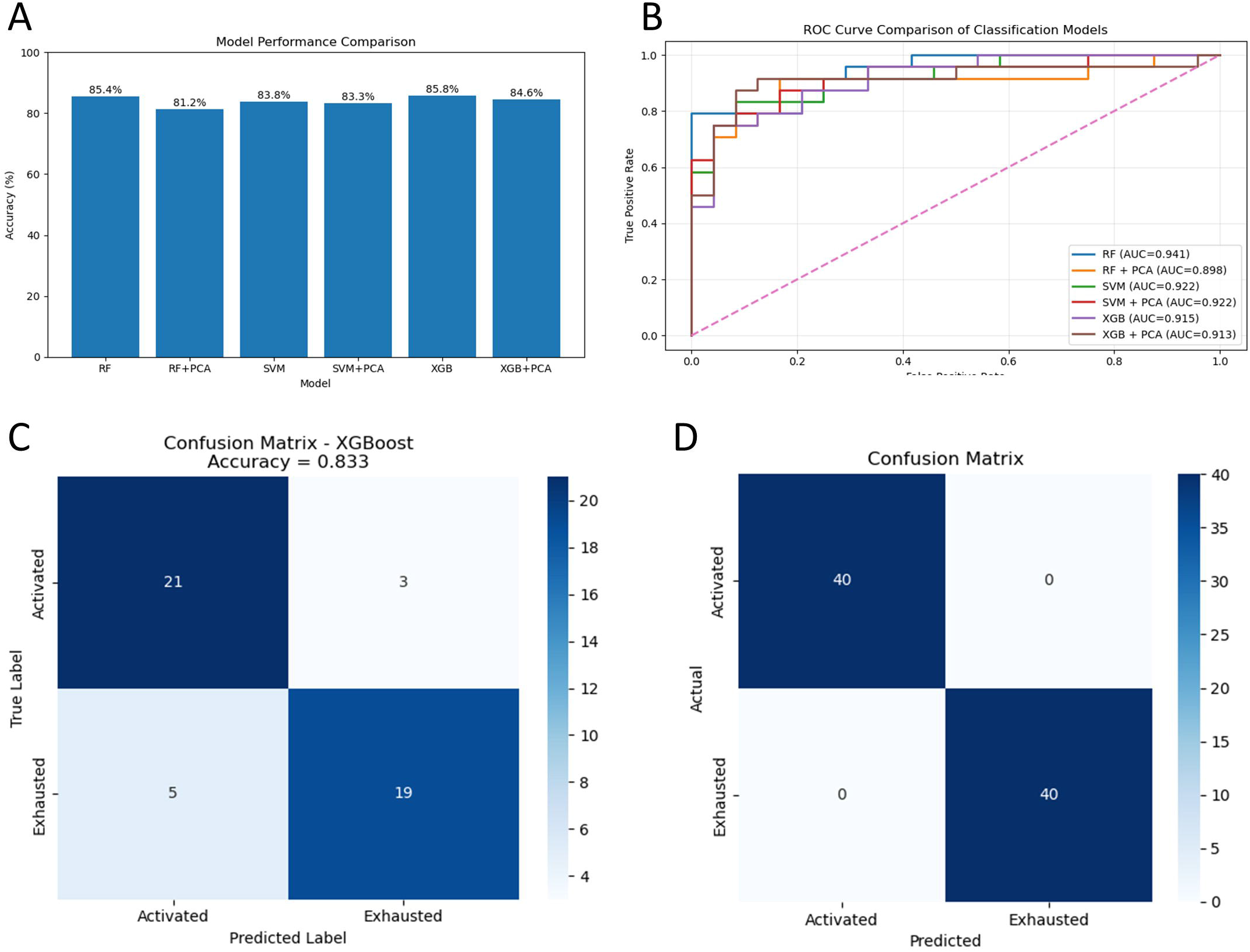
Machine learning-based classification of activated and exhausted CD8^+^ T cells using Raman spectral signatures. (A) Comparison of classification accuracy among different machine learning models (B) ROC analysis of machine learning models for T-cell state classification (C) Confusion matrix obtained from the XGBoost classifier for activated and exhausted T-cell classification (D) External validation of the XGBoost classifier using independent biological samples.

## Discussion

T cell exhaustion remains a major barrier to effective cancer immunotherapy, as dysfunctional CD8^+^ T cells fail to sustain durable anti-tumor responses within the tumor microenvironment. Therefore, a deeper understanding of exhausted T cells and their accurate identification within tumors is essential for improving therapeutic intervention and patient prognosis. While exhausted T cells can be studied directly from tumors, the low yield of antigen-specific TILs, cellular heterogeneity, and the complex influence of the tumor microenvironment often complicate mechanistic investigations. In contrast, in vitro systems offer a controlled, reproducible, and readily accessible platform for inducing and tracking exhaustion over time.

To this end, we established an in vitro model of CD8^+^ T cell exhaustion through chronic antigenic stimulation. This model recapitulated key phenotypic, functional, transcriptional, and metabolic features of exhausted T cells. Progressive loss of proliferation, pro-inflammatory cytokine secretion, and cytotoxicity, along with altered Ly108 expression dynamics representing different stages of exhaustion from progenitor to terminal states, suggested a gradual transition of cells towards terminal exhaustion. Following the establishment of the progressive exhaustion model, chronically stimulated day 6 (CS6) cells were identified as bona fide exhausted CD8^+^ T cells and were further characterized for additional exhaustion-associated markers.

Although CS6 cells exhibited higher expression of inhibitory receptors such as PD-1 and Lag3 compared to AS6 cells, variability across markers including TIGIT, TIM-3, and CTLA-4 highlighted the heterogeneous and dynamic nature of exhaustion-associated phenotypes. However, when co-expression of inhibitory receptors were considered, CS6 cells showed clusters in t-SNE plots, which were absent in AS cells thereby supporting the acquisition of an exhaustion-associated phenotype.

Apart from inhibitory receptor expression and functional decline, exhausted T cells undergo profound metabolic adaptations within the TME, including altered lipid metabolism and impaired bioenergetic fitness. Consistent with this, CS6 cells displayed elevated expression of the lipid uptake receptor CD36 and the ecto-nucleotidase CD39. Increased CD39 expression further supported the acquisition of a terminally differentiated phenotype. CD39 is frequently co-expressed alongside inhibitory receptors on exhausted T cells and hydrolyzes extracellular ATP into AMP, which is subsequently converted into immunosuppressive adenosine by CD73. Adenosine suppresses T cell effector functions and contributes to immune dysfunction within the TME. In parallel, exhaustion-associated transcriptional rewiring was also observed. TCF-1, which sustains stemness and progenitor-like functionality in T cells, was markedly downregulated in CS6 cells. In contrast, TOX expression was elevated, supporting the establishment and maintenance of the exhausted phenotype. TOX not only promotes inhibitory receptor expression but also drives long-term epigenetic remodeling associated with T cell exhaustion.

Following the characterization of bona fide exhausted cells, Raman spectral signatures of exhausted, and activated, CD8^+^ T cells were acquired and compared. Raman spectral profiling revealed distinct biochemical differences among the two cellular states. Activated cells exhibited enhanced nucleic acid-, protein-, and lipid-associated spectral features, consistent with elevated metabolic and biosynthetic activity. In contrast, exhausted cells displayed comparatively reduced spectral intensities and localized in a different plane from the activated cells in the PCA space. A deeper analysis of the Raman spectra through AUC quantification of biomolecule-associated bands revealed distinct biochemical features underlying the different T cell states. Importantly, many of the spectral changes observed in exhausted cells aligned with known metabolic and molecular adaptations reported during T cell exhaustion. The concordance between Raman-derived biochemical signatures and established exhaustion-associated pathways highlights the potential of Raman spectroscopy as a label-free approach for identifying and characterizing exhausted CD8^+^ T cells.

Beyond unsupervised dimensionality reduction using PCA, the spectral differences identified by Raman spectroscopy were distinct enough to classify T cell states reliably, using supervised machine learning approaches. Among the models tested, the XGBoost pipeline achieved the highest accuracy with minimal variation across validation runs, demonstrating that the biochemical signatures captured by Raman spectroscopy can be leveraged for robust identification of exhausted CD8^+^ T cells.

Collectively, this study demonstrates that label-free single-cell Raman spectroscopy can capture biochemical alterations associated with CD8^+^ T cell exhaustion and distinguish exhausted cells from activated counterparts without extensive sample processing. The distinct spectral signatures observed in exhausted cells reflected alterations in cellular functionality and metabolic state, particularly in comparison to activated cells. The ability of Raman spectroscopy to resolve functional T cell states based on intrinsic biochemical composition highlights its potential as a platform for immune profiling and monitoring responses to immunotherapeutic interventions, including checkpoint blockade and metabolic reprogramming strategies. Furthermore, coupling Raman spectroscopy with machine learning algorithms may provide a rapid, label-free platform for the identification and quantification of exhausted T cells in clinical samples. Given that the abundance and severity of T cell exhaustion have been linked to patient prognosis and therapeutic responsiveness, such an approach could have utility as a prognostic modality and clinical decision-support tool. The ability to objectively assess exhaustion-associated biochemical signatures may also aid patient stratification and monitoring of responses to immunotherapeutic interventions. Together, these findings establish Raman spectroscopy as a promising tool for real-time characterization of T cell exhaustion in cancer and chronic infections.

## Limitations

1. The Raman bands identified in this study were assigned based on existing literature; however, several of the discriminatory peaks lie within complex spectral regions where contributions from multiple biomolecules overlap. Therefore, further peak deconvolution and detailed spectral analyses will be necessary to identify the specific molecular vibrations responsible for the observed differences between activated and exhausted T cells.
2. In addition, the study was performed using murine OT-1 CD8^+^ T cells, which provide a controlled and reproducible model for investigating T cell exhaustion. While this system was suitable for establishing exhaustion-associated Raman signatures, translation of these findings to a clinical setting will require validation in human exhausted CD8^+^ T cells, particularly from patient-derived samples.

## Acknowledgements

The authors thank Debashree Guha Adhya for critical inputs in the ML-based classification models. AAB, PA, PSM, acknowledge Ministry of Education (MoE), Govt. of India, for fellowship. SB acknowledges DBT for fellowship. The authors also acknowledge Central Research Facility of IIT Kharagpur for SPR facilities. ChatGPT was used as a paraphrasing tool in the manuscript.

## Author contributions

A.A.B.: Designing and performing research, data analysis, writing original draft; P.A..: Raman spectroscopy and analysis; P.S.M.: PCA analysis; A.T..: performing ML-based classification of spectral data; S.B.: In vitro experiments and data analysis; S.D.: Supervised Raman spectroscopy and analysis, G.M.: Conceptualization and designing research, analysis, and supervision, writing of manuscript.

## Ethical approval statement

All experiments using mice were executed in agreement with guidelines established by the Institutional Animal Ethical Committee, IIT Kharagpur (Ethical certificate no. IE-09/GM-SMST/1.22).

